# The gene expression profile of the song control nucleus HVC shows sex specificity, hormone responsiveness, and species specificity among songbirds

**DOI:** 10.1101/2021.02.28.432133

**Authors:** Meng-Ching Ko, Carolina Frankl-Vilches, Antje Bakker, Manfred Gahr

## Abstract

Singing occurs in songbirds of both sexes, but some species show typical degrees of sex-specific performance. We studied the transcriptional sex differences in the HVC, a brain nucleus critical for song pattern generation, of the forest weaver (*Ploceus bicolor*), the blue-capped cordon-bleu (*Uraeginthus cyanocephalus*), and the canary (*Serinus canaria*), which are species that show low, medium, and high levels of sex-specific singing, respectively. We observed persistent sex differences in gene expression levels regardless of the species-specific sexual singing phenotypes. We further studied the HVC transcriptomes of defined phenotypes of canary, known for its testosterone-sensitive seasonal singing. By studying both sexes of canaries during both breeding and nonbreeding seasons, nonbreeding canaries treated with testosterone, and spontaneously singing females, we found that the circulating androgen levels and sex were the predominant variables associated with the variations in the HVC transcriptomes. The comparison of natural singing with testosterone-induced singing in canaries of the same sex revealed considerable differences in the HVC transcriptomes. Strong transcriptional changes in the HVC were detected during the transition from nonsinging to singing in canaries of both sexes. Although the sex-specific genes of singing females shared little resemblance with those of males, our analysis showed potential functional convergences. Thus, male and female songbirds achieve comparable singing behaviours with sex-specific transcriptomes.

## Introduction

Most of the genomes of male and female individuals of the same species are the same but stark sex differences in physiological, phenotypical, or behavioural traits between the sexes are common and widespread in the animal kingdom. The songbird clade (a suborder of the perching birds) consists of more than 4000 extant avian species, and these exhibit a great diversity of sex differences in singing behaviour. Among domesticated canaries (*Serinus canaria*), males are known for their singing behaviour and have been selected for their sophisticated songs for centuries, whereas female canaries seldom sing (Hartley et al., 1997; Herrick and Harris, 1957; Ko et al., 2020; Pesch and Güttinger, 1985; Shoemaker, 1939; Vallet et al., 1996). Such substantial sex differences in singing behaviour are commonly found in the majority of northern temperate songbird species, even though females of many tropical and southern temperate species sing regularly, and their songs play an important role in inter-sexual communication (Hall et al., 2015; Price, 2019; Price et al., 2009; Slater and Mann, 2004). For example, female blue-capped cordon-bleus (*Uraeginthus cyanocephalus*) use their song as advertising signals and address their songs to their mates (Immelmann, 1968; Ota et al., 2018), although the female songs appear shorter and less complex than those of males (Geberzahn and Gahr, 2011). In another tropical songbird species, forest weavers (*Ploceus bicolor*), males and females develop their songs during pair binding and eventually learn to sing identical duets, which they use to defend their territories (Wickler and Seibt, 1980).

Although the extent of sex differences in singing behaviour varies greatly in the songbird clade (Ball, 2016), the song quality and occurrence in males and females of many songbird species are dependent on testosterone (Dittrich et al., 2014; Fusani et al., 2003; Ko et al., 2020; Leitner et al., 2001b; Nottebohm et al., 1987; Voigt and Leitner, 2008). In canaries that breed seasonally, the optimal breeding conditions are tightly associated with an increase in the day length, which initiates gonadal growth and testosterone production as well as various types of breeding activity, including singing (Leitner et al., 2001b; Nottebohm et al., 1987; Voigt and Leitner, 2008). The length and syllable repetition rate of songs during the breeding season are greater than those of songs in nonbreeding seasons (Leitner et al., 2001a; Leitner et al., 2001b; Voigt and Leitner, 2008). Castrated male canaries sing shorter songs than sham-operated males, and the subcutaneous implantation of testosterone results in the recovery of singing performance (Heid et al., 1985). The local administration of testosterone into the preoptic brain region increases the male canary singing rate by increasing motivation (Alward et al., 2013). Similarly, the systemic administration of testosterone reliably and repeatedly induces singing behaviour in otherwise nonsinging female canaries (Fusani et al., 2003; Herrick and Harris, 1957; Leonard, 1939; Madison et al., 2015; Nottebohm, 1980; Shoemaker, 1939; Vellema et al., 2019b). Although female canaries rarely exhibit spontaneous singing (Hartley et al., 1997; Herrick and Harris, 1957; Ko et al., 2020; Pesch and Güttinger, 1985; Shoemaker, 1939; Vallet et al., 1996), its occurrence appears to be associated with the plasma androgen levels (Ko et al., 2020).

A set of interconnected neural circuits called the song control system controls the production and learning of singing behaviour (Nottebohm et al., 1976; Wild, 2004). The premotor nucleus HVC (used as the proper name), which is a sensorimotor integration centre in the song control system, is involved in the frequency and temporal modulation of songs in male and female songbirds (Hahnloser et al., 2002; Hoffmann et al., 2019) and in the sexual preferences regarding conspecific song displays (Brenowitz, 1991; Del Negro et al., 1998). The HVC is the only brain nucleus within the song control system that expresses receptors for both androgens and estrogens (Frankl-Vilches and Gahr, 2018; Gahr, 2001). Intriguingly, the anatomical properties of the HVC, such as volume (Figure 1), neuron number, and dendrite complexity, are male-biased (greater, higher and more complex, respectively, in males than in females) in all songbird species that have been examined, irrespective of the existence of sex differences in singing behaviour (Brenowitz et al., 1985; Gahr et al., 2008; Gahr et al., 1998; Gurney and Konishi, 1980; Hall et al., 2010; Lobato et al., 2015; MacDougall-Shackleton and Ball, 1999; Nixdorf et al., 1989; Nottebohm and Arnold, 1976; Schwabl et al., 2015). Testosterone treatment increases the delineable volume of the HVC in both male and female canaries (Fusani et al., 2003; Madison et al., 2015; Nottebohm, 1980) and many other species (Bernard and Ball, 1997; Dittrich et al., 2014; Gulledge and Deviche, 1999; Smith et al., 1997; Van Meir et al., 2004). Nevertheless, the HVC volume of female canaries implanted with testosterone remains markedly smaller than that of males (Nottebohm, 1980) (Figure 1). Testosterone clearly regulates singing behaviour and the anatomy of the HVC in both male and female songbirds. However, an intrinsic limit to the alterations induced by testosterone appears to exist, and this limit prevents female songbirds from reaching the same levels of “maleness” in terms of song characteristics and HVC anatomy.

**Figure 1.**
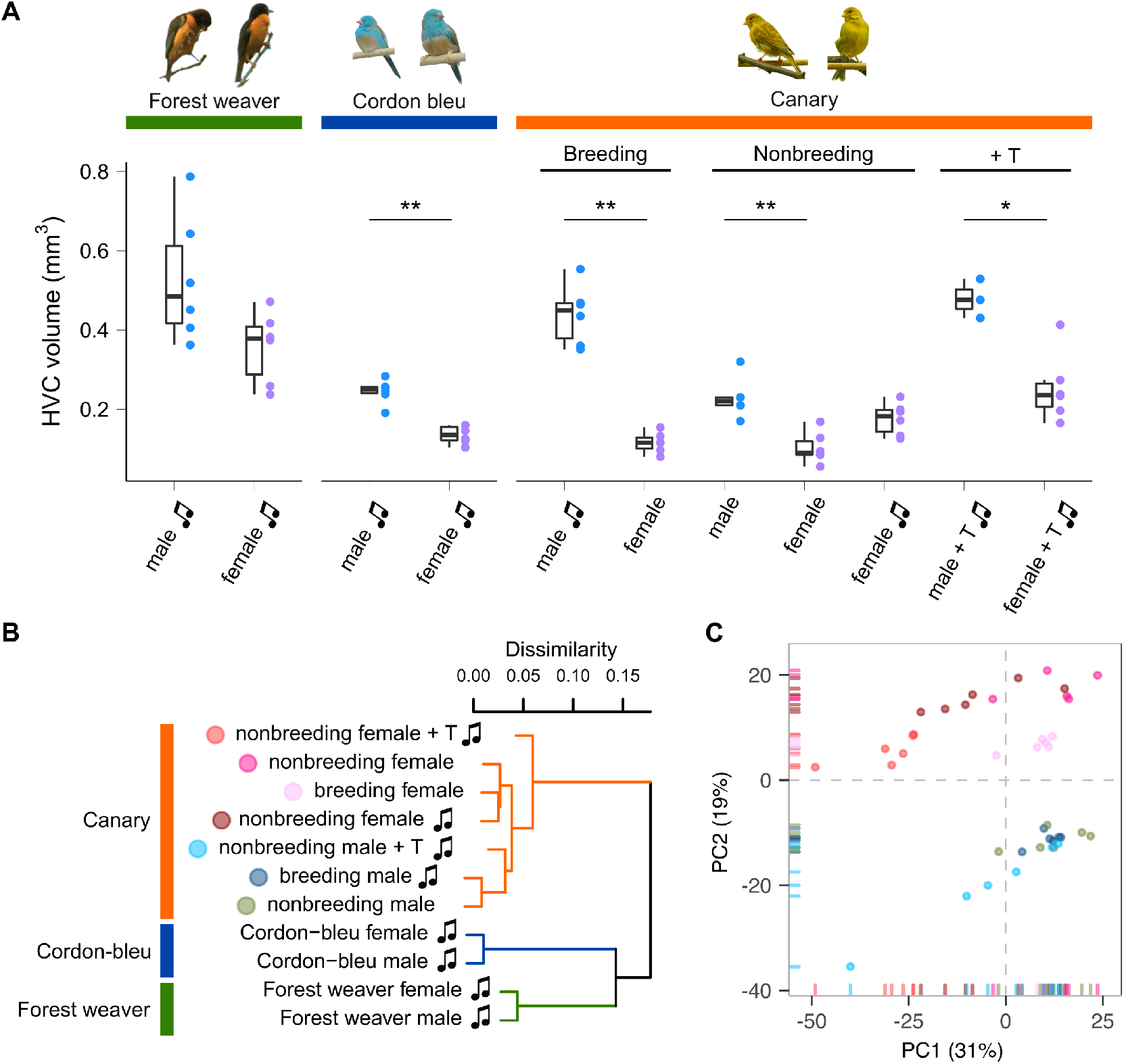
Species, sex, and plasma androgen level are the major determinants of the HVC gene expression patterns in songbirds. **A**, HVC volume of birds used in this study. Forest weavers: males 0.528 mm^3^ (mean), females 0.357 mm^3^. Mann-Whitney Test, U = 7, P value = 0.0931. Cordon-bleus: males 0.245 mm^3^, females 0.136 mm^3^. Mann-Whitney Test, U = 0, P value = 0.00217. Breeding canaries: singing males 0.439 mm^3^, nonsinging females 0.116 mm^3^. Mann-Whitney Test, U = 0, P value = 0.00217. Nonbreeding canaries: nonsinging males 0.228 mm^3^, nonsinging females 0.103 mm^3^, singing females 0.176 mm^3^. Mann-Whitney Test (nonsinging males vs. nonsinging females), U = 0, P value = 0.00492. Mann-Whitney Test (nonsinging males vs. singing females), U = 8, P value = 0.127. Testosterone-implanted canaries: males 0.478 mm^3^, females 0.253 mm^3^. Mann-Whitney Test, U = 0, P value = 0.0238. * P value < 0.05; ** P value < 0.01. Each colour-coded dot indicates the measurement from one bird. The boxes indicate the 25th/50th/75th percentiles (bottom/middle/top bar), and the extent of the whiskers indicates the most extreme values that are within 1.5 times the IQR (interquartile range) of the hinge. T: testosterone; music note: presence of singing behaviour. **B**, Hierarchical clustering showing that phylogenetic relatedness accounts for the most variation in the HVC transcriptomes of females and males of the forest weaver, cordon-bleu, and canary. Among the seven canary groups, the testosterone-treated animals were the least similar to the untreated canaries. **C**, A PCA of the HVC transcriptomes of the seven canary groups distinguished male birds (PC2 < 0) from female birds (PC2 > 0). Each point is colour-coded by group (see B) and represents an HVC sample.

How are such sex-specific differences in singing and HVC anatomy achieved in different species? We hypothesize the existence of a fundamental difference in gene regulation and expression between male and female conspecific songbirds. Based on this hypothesis, sex differences in the HVC transcriptomes of male and female conspecifics should always be observable, even though both sexes share an almost identical genome. The female-specific W chromosome harbours 30-50 genes, whereas one and two copies of the Z chromosome (harbouring approximately 1,080 genes) are present in female and male birds, respectively (Frankl-Vilches et al., 2015; Sayers et al., 2020; Smeds et al., 2015). We further hypothesize that such intrinsic differences cannot be overwritten by manipulating the testosterone level in females. To test our hypotheses, we compared the transcriptomes of HVCs microdissected from three songbird species—the forest weaver, the blue-capped cordon-bleu, and the canary. These species represent three categories of sex differences in singing behaviour (low, medium, and high). Furthermore, we compared male and female canaries with and without testosterone implantation to evaluate the effects of testosterone (Table 1). The results revealed sex differences in the HVC transcriptome between- and within-species comparisons. Our results indicate that although the extent of sex-biased gene expression is context-dependent, it is persistent regardless of hormonal manipulation, behavioural phenotypes, and distinct genetic backgrounds.

**Table 1.**
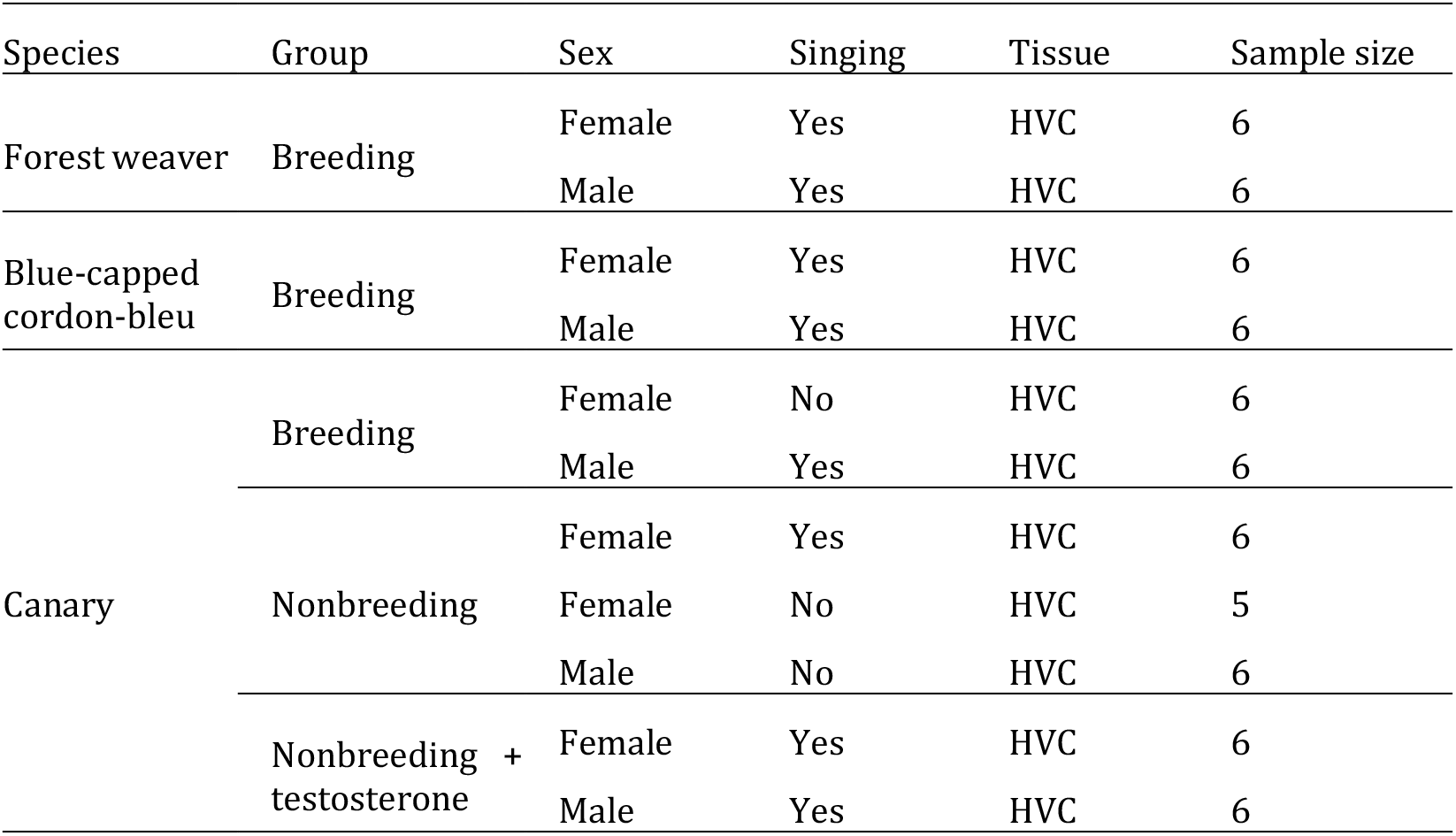
Experimental groups and sample sizes.

## Materials and methods

### Animals

Forest weaver (*Ploceus bicolor*) pairs were observed in their breeding territories in eastern South Africa; the songs of the pairs were recorded to ensure that both mates were singing. Subsequently, the animals were caught and sacrificed in accordance with permits issued by the local authorities (Chief Professional Officer for Research at the Natal Parks, Game and Fish Preservation Board, P. O. B. 662, Pietermaritzburg 3200). Blue-capped cordon-bleus (*Uraeginthus cyanocephalus*) and canaries (*Serinus canaria*) were bred at the animal facilities at the Max Planck Institute for Ornithology in Seewiesen, Germany. The procedures used for animal housing and welfare complied with the European directives for the protection of animals used for scientific purposes (2010/63/EU), and the protocols were approved by the Government of Upper Bavaria (AZ No. 55.2-1-54-2532-181-12). Adult canaries and blue-capped cordon-bleus (aged at least 1 year) were housed in pairs under long-day conditions (light:dark = 14:10 hours), and their breeding activities were monitored. The birds (male canaries and cordon-bleus) were sacrificed after singing activity was observed during the breeding season. Nonbreeding canaries were housed pairwise with a 9-hour light:15-hour dark schedule for at least 8 weeks before their singing activity was monitored (generally starting in late September). Nonbreeding male and female canaries were maintained alone in sound-attenuated boxes (70 × 50 × 50 cm), recorded continuously during the song monitoring phase (two weeks) and sacrificed after confirmation of no singing activity. Additional groups of male and female nonbreeding and nonsinging canaries were implanted with testosterone for 2 weeks and sacrificed after observation of singing activity. Water and food were available *ad libitum*. The sex was confirmed by PCR using the P2 and P8 primers for CHD genes (Griffiths et al., 1998) and by visual inspection of the reproduction system after sacrifice. We previously found that six nonbreeding female canaries exhibited singing behaviour during long-term monitoring, and their songs have been described in detail (Ko et al., 2020). In this study, we included the transcriptomes of these birds. All the birds were sacrificed via an overdose of isoflurane followed by decapitation, the body weights were recorded, and the brains were dissected, weighed, snap-frozen on dry ice and stored at - 80°C until further use. Table 1 summarizes the information of the experimental groups used in this study.

### Testosterone implantation

A Silastic™ tube (Dow Corning; 1.47-mm inner diameter, 1.96-mm outer diameter, 0.23-mm thickness) was cut to a length of 7 mm and loaded with testosterone (86500, Fluka) as densely as possible. The two ends of the Silastic™ tube were sealed with silicone elastomer (3140, Dow Corning). After closure, the implants were cleaned with 100% ethanol to remove testosterone particles and then immersed in ethanol overnight in a hood to ensure no leakage at either end. Implants with apparent dampness were discarded. One day prior to the implantation, the implants were incubated in 0.1 M phosphate buffered saline (PBS) overnight to enable the immediate release of testosterone upon implantation (Rasika et al., 1994). We started implantation at approximately 8:30 am (immediately after the light was turned on in the morning), which resulted in a 20-minute interval between birds based on the scarification time. A small incision was made on the back of the bird over the pectoral musculature, and one testosterone implant was placed subcutaneously. The skin was closed by the application of tissue glue. After 2 weeks, the animals were sacrificed. The testosterone implants were checked, and this inspection revealed that the implants were all in place and were not empty at the end of the experiments.

### Radioimmunoassay of plasma testosterone

Blood was sampled (< 150 μl) at the time sacrifice. All blood samples were taken between 8 and 11 am and were taken within 3 min after caught to avoid the effect of handling (Wingfield et al., 1982). Blood samples were centrifuged (5,000 rpm, 10 min) to separate the plasma from blood cells. Testosterone metabolites were measured with a radioimmunoassay using a commercial antiserum against testosterone (T3-125, Endocrine Sciences, Tarzana, USA) as previously described (Goymann et al., 2002). Standard curves and sample concentrations were calculated with Immunofit 3.0 (Beckman Inc. Fullerton, CA) using a four-parameter logistic curve fit and corrected for individual recoveries.

The testosterone concentrations were assayed in duplicate in five separate assays. The mean extraction efficiency for plasma testosterone was 85.0 ± 3.9% (mean ± SD, N = 42). The lower detection limits of the testosterone assays were 0.34, 0,35, 0.36, 0.38 and 0.35 pg per tube, and all the samples were above the detection limit. The intra-assay coefficients of variation of a chicken plasma pool were 8.7%, 3.4%, 12.8%, 1.9%, and 4.4%. The interassay coefficient of variation as determined by the variation in the chicken plasma pool between all the assays was 5.1%. Because the testosterone antibody used shows significant cross-reactions with 5α-dihydrotestosterone (44%), our measurement might include a fraction of 5α-DHT.

### Brain sectioning

The birds were killed by an overdose of isoflurane, and their brains were snap-frozen on dry ice. The brains were sectioned sagittally into four series of 40-μm sections and two series of 20-μm sections with a cryostat (Jung CM3000 Leica). The 40-μm sections were mounted on glass slides for subsequent tissue dissection for total RNA extraction, whereas the thin sections were mounted on RNase-free Superfrost slides for Nissl staining and measurement of the HVC volume. All sections were stored at -80°C until further processing.

### Measurement of the HVC volume

One series of 20-μm sections mounted on RNase-free Superfrost slides was subjected to Nissl staining with 0.1% thionin (Sigma-Aldrich), dehydrated, immersed in xylene and cover-slipped with Roti-Histokitt II mounting medium (Roth). The HVC areas (typically 8-10 slices) were measured with a Leica DM6000 B microscope connected to a computer-based image-analysis system (IMAtec). All brains were coded such that the observers were blind towards any additional information about the sections they measured during the delineations. The volumes were derived from the summed area measurements multiplied by the section thickness and the intersection distance.

### Microarray procedures and annotation

For total RNA extraction, the song control nucleus HVC and the visual area of the entopallium were dissected from the abovementioned 40-μm sections under a stereomicroscope (typically 24-32 slices for the HVC and 16-20 slices for the entopallium) and transferred into an Eppendorf tube containing 340 μl of RLT buffer mixture (containing DTT, Qiagen). This dissection procedure using rather thin sections reduces the contamination of HVC tissue with surrounding tissue. RNA was then extracted using the RNeasy® Micro Kit (Qiagen) with the optional DNase digest step. The RNA quality was assessed using the Agilent Model 2100 Bioanalyzer (Agilent Technologies), and the RNA concentrations were assessed using a Nanodrop 1000 spectrometer (Thermo Fisher Scientific). The RNA quality of all samples was good (RIN > 7). The purified total RNA samples (at least 100 ng per sample) were subsequently processed and hybridized using the Ambion WT Expression Kit and the Affymetrix WT Terminal Labelling and Controls Kit. The resulting cDNA was hybridized to the Custom Affymetrix Gene Chip® MPIO-ZF1s520811 Exon Array, which has been used successfully and validated in cross-species hybridization studies (Dittrich et al., 2014; Frankl-Vilches et al., 2015). The 5.7 million male zebra finch-specific probes spotted on this array correspond to approximately 4,711,133 probe sets and hence to 25,816 transcripts published in public databases (NCBI and Ensembl) (Warren et al., 2010). We annotated more than 90% of the transcripts to 12,729 human orthologous genes using several publicly available databases (Ensembl, GenBank, UniProt, and DAVID (Benson et al., 2005; Consortium, 2015; Flicek et al., 2014; Huang et al., 2008, 2009; Yates et al., 2016)) and commercial databases (El Dorado, Genomatix, Precigen Bioinformatics Germany GmbH (PBG), Munich, Germany). Hybridization was performed for 16 hours at 45°C and 60 rpm in a GeneChip Hybridization Oven 640 (Affymetrix). The arrays were washed, stained, and scanned using the Affymetrix GeneChip Fluidics Station 450 and the Affymetrix GeneChip scanner 3000 7G. The CEL files were generated using Affymetrix® GeneChip® Command Console® Software (AGCC), and for quality control, the success of individual hybridizations was assessed using Affymetrix® Expression Console™ software.

Differential expression was calculated using ChipInspector software version 21 (El Dorado Database version E28R1306 (Genomatix)). ChipInspector is a single probe-based analysis tool for microarray data that can show increased sensitivity compared with that obtained with conventional probe set-based analyses, such as robust multiarray analysis. ChipInspector consists of four steps: single probe-transcript annotation (ensuring up-to-date annotation), total intensity normalization, SAM (significance analysis of microarrays, adapted to single-probe handling) analysis (Tusher et al., 2001), and transcript identification based on significantly changed probes (Cohen et al., 2008). We set the delta threshold to 0 (to control the false positive rate) and used the groupwise exhaustive comparison tool in ChipInspector. The minimum coverage for each transcript was set to 10 significant probes, and the minimum expression difference was |log2(fold change)| ≥ 0.5. The significantly differentially expressed transcripts obtained were annotated to human orthologous genes as described above. For transcripts belonging to the same genes, the average expression was calculated if all transcripts were regulated in the same manner (e.g., all upregulated or all downregulated). If a gene contains both up- or downregulated transcripts, the transcripts that showed changes in expression in the minority direction were discarded (<40% of the total transcripts), and the average expression was instead calculated from the remaining transcripts (60% or more). Transcripts without human orthologous gene annotation were removed prior to subsequent analyses. We were cautious about possible cross-species bias in hybridization, and differential expression analyses were only performed between two conspecific groups. The microarray data discussed in this publication have been deposited in NCBI’s Gene Expression Omnibus (Edgar et al., 2002) and are accessible through the GEO Series accession number <u>GSE83674</u>.

### Hierarchical clustering

The normalized gene expression levels (across all transcriptome samples) were calculated using the “justRMA” function of the R package “affy” v1.66.0 (Gautier et al., 2004), and the expression levels were further collapsed to gene levels using the “collapseRows” functions of the R package “WGCNA” v1.69 (Langfelder and Horvath, 2008, 2012). The group-level expression levels were calculated by averaging the gene expression levels. Spearman’s ρ was calculated between samples (or groups) using the “cor” function, and a Euclidean distance matrix was calculated (dist(1-cor)) and used for hierarchical clustering analysis (method: complete) with the “hclust” function. This computation was performed with R v4.0.2 (R_Core_Team, 2020) using the “dendextend” v1.14.0 package for visualization (Galili, 2015).

### Principal component analysis (PCA)

PCA (Ringner, 2008) was performed using the normalized and collapsed HVC gene expression data (described in the hierarchical clustering section) from 41 canaries (Table 1). The computation was performed using the “pca” function of the R package “pcaMethods” (v1.60.0 (Stacklies et al., 2007), method = svd). The data were centred but not scaled because the expression data had already been normalized.

### Fisher’s exact testing for chromosomal enrichment

Fisher’s exact test was used to test whether sex-biased genes were enriched on a chromosome by comparing the gene lists of interest to the zebra finch annotation (https://www.ncbi.nlm.nih.gov/genome/?term=zebra+finch). We used the “fisher.test” function in R (R_Core_Team, 2020) with the alternative set to “greater” for enrichment. The male- and female-biased genes identified from each comparison as well as the male-specific, female-specific, and sex-shared genes were tested separately. The P values were adjusted using the Bonferroni correction to account for multiple comparisons using “p.adjust” in R (R_Core_Team, 2020).

### Gene Ontology (GO) term enrichment analysis

We used ClueGO v2.2.4, an application built in the Cytoscape environment v3.3.0 (Bindea et al., 2009; Shannon et al., 2003), to predict the putative biological functions of the genes of interest. This software performs GO term enrichment hierarchical analyses and fuses GO terms with similar functions. The enrichment was determined by the right-sided test and corrected using the Bonferroni step-down method considering multiple comparisons.

## Results and discussion

### The species identity distinguishes its HVC transcriptomes

We quantified the total mRNA from the HVC of male and female birds belonging to three songbird species (forest weavers, blue-capped cordon-bleus, and canaries) during the breeding season (Table 1). For intraspecies sex comparisons, we included male and female canaries during the nonbreeding seasons (both nonsinging), and subgroups were treated with testosterone to induce singing behaviour in both sexes (Supplementary Figure 1). In addition, we obtained a rare group of nonbreeding female canaries that sang spontaneously without exogenous testosterone manipulation. However, their plasma androgen levels were intrinsically higher than those of nonbreeding and nonsinging females (Ko et al., 2020). To visualize the similarities in the gene expression patterns among and within the three species, we calculated Spearman’s rank correlation coefficient σ for 12,360 gene expression levels and calculated the distance matrix based on the coefficient σ. The resulting cladogram indicated that the HVC transcriptomes of the eleven groups were clustered primarily by phylogenetic relatedness (Figure 1B and Supplementary Figure 2). We obtained a similar result with the transcriptomes of another tissue, the entopallium (Supplementary Figure 3), which is an area of the avian visual system that is functionally equivalent to the primary visual cortex of mammals. The within-species comparison of the canary HVC showed that the individual transcriptomes clustered well by sex, with the exception of the nonbreeding canaries implanted with testosterone (Figure 1B and Supplementary Figure 2). The subclusters of forest weaver and cordon-bleu HVC transcriptomes were less sexually differential than those of canaries, i.e., male and female birds were intermittent within the subcluster of the two species (Supplementary Figure 2 and Supplementary Figure 3).

### Sex-biased gene expression in the HVC of songbirds

We defined the degree of sex differences in HVC gene expression levels as the number of sex-biased genes whose expression levels showed significant differences between males and females of the same species (male-biased genes showed higher expression levels in males than in females, log_2_ (fold change) ≥ 0.5; female-biased genes presented higher expression levels in females than in males, log_2_ (fold change) ≤ -0.5). For canary, we quantified the sex differences from five distinct male-to-female comparisons (Figure 2A, comparisons 3 to 7).

**Figure 2.**
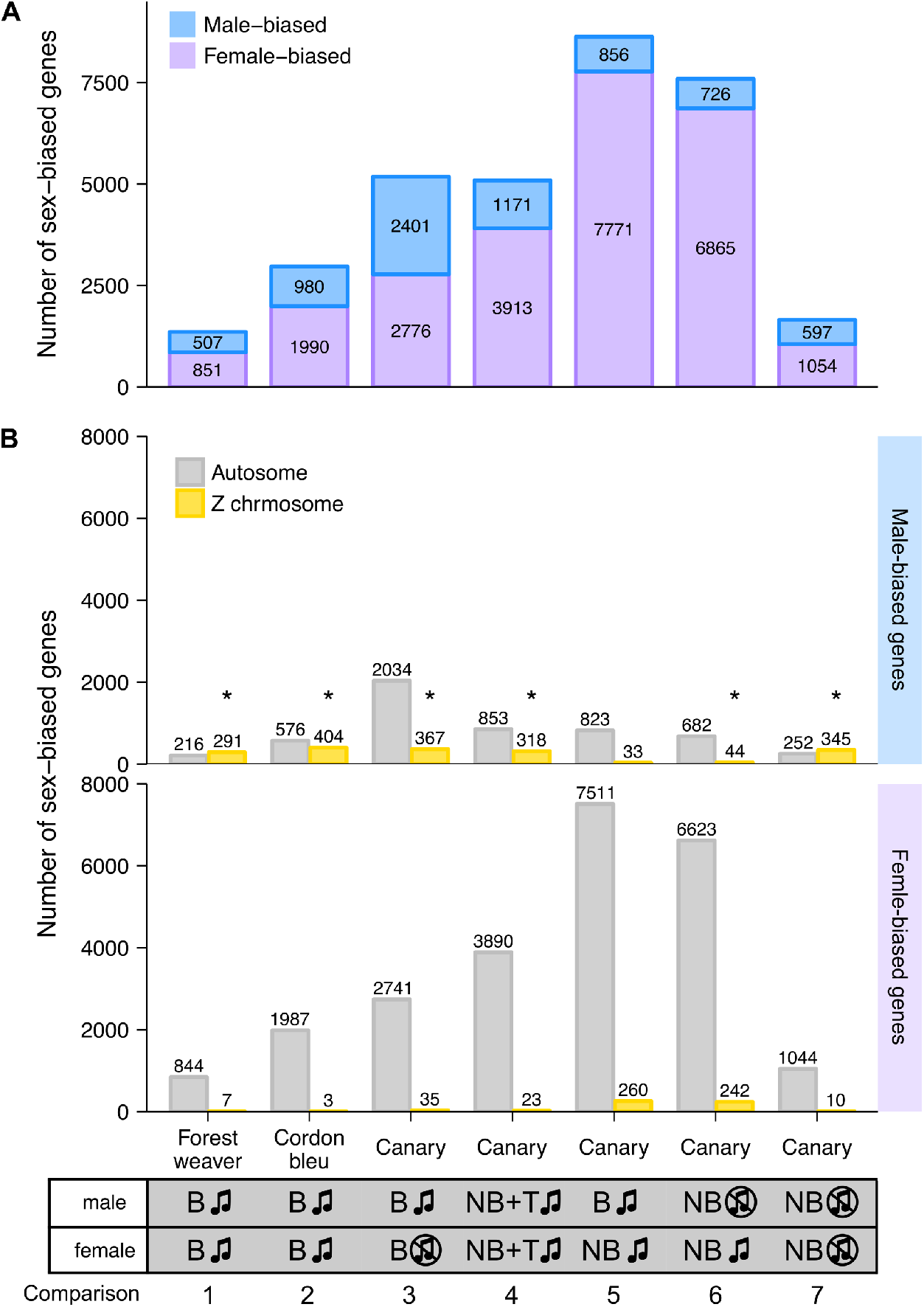
Female- and male-biased sex differences in HVC gene expression levels in songbirds. **A**, The bar graph summarizes the number of sex-biased genes in the HVC transcriptome identified from each male-to-female comparisons. The phenotypes of the groups being compared in each male-to-female comparison are listed at the bottom of the graph. **B**, The bar graph summarizes the number of autosomal and Z chromosomal genes classified as male- and female-biased genes in the HVC transcriptome based on each male-to-female comparison. Abbreviations: B, breeding; NB, nonbreeding; T, testosterone implantation. The music note indicates the presence or absence of singing behaviour; the asterisk (*) indicates enrichment for Z chromosomal genes (adjusted p < 0.05, Fisher’s exact test followed by the Bonferroni correction for multiple testing).

Interestingly, we found persistent sex differences in HVC gene expression levels from all the comparisons, regardless of the degree of sex differences in singing behaviour (Figure 2A). The majority (> 90%) of sex-biased genes were lowly biased (0.5 ≤ |log2 (fold change)| < 1, Supplementary Figure 4). The comparison of forest weaver (the female birds can sing identical songs to the males; least differences in the HVC volumes between the sexes; Figure 1A) showed the least prominent sex differences in the HVC transcriptomes across all the comparisons performed in this study. Nevertheless, the extent of sex differences in gene expression levels was extensive (> 1,300 sex-biased genes, Figure 2A, comparison 1). The comparison showing the second-lowest degree of sex-differences was the comparison of nonbreeding male and nonbreeding female canaries (> 1,500 sex-biased genes, Figure 2A, comparison 7), both lacking singing behaviour, but HVC volume was male biased (Figure 1A). Testosterone implantation induced singing behaviour and increased the HVC volume of both male and female canaries, although a markedly smaller HVC volume continued to be observed in these females (Figure 1A and Supplementary Figure 1C). Such processes in females are referred to as “masculinization” of the female brain and behaviour (Arnold and Gorski, 1984; Wade, 2001). However, the gene expression levels of the testosterone-stimulated singing female canaries were very different from those of the male canaries administered the same treatment (> 5,000 sex-biased genes, Figure 2A, comparison 4). The comparisons showing the most striking sex differences were the two that included spontaneously singing female canaries. As revealed in these comparisons, many genes showed markedly different regulation in spontaneously singing female canaries, as reflected by expression levels, compared with breeding singing male canaries (> 8,600 sex-biased genes, comparison 5) and nonbreeding nonsinging male canaries (> 7,500 sex-biased genes, comparison 6, Figure 2A).

For each comparison, we decomposed the sex-biased genes into two classes, autosomal genes and Z sex chromosomal genes, to assess whether the sex-biased genes were mainly concentrated on the sex chromosomes. Annotation for W chromosomal genes was unfortunately not available in our study. We used Fisher’s exact test to evaluate whether a particular chromosome was enriched in sex-biased genes. We observed enrichments of Z chromosomal genes among the male-biased genes identified in most of the comparisons (Figure 2B and Supplementary Table 1). This observation was not surprising because males are the homogametic sex (have two copies of Z), whereas females are the heterogametic sex (have one copy of Z and one copy of W) in birds, and the dosage compensation for Z-linked genes is less complete in birds than that in mammals (Itoh et al., 2007; Mank, 2013; Nätt et al., 2014; Uebbing et al., 2015; Wolf and Bryk, 2011). However, one canary comparison (breeding singing males and nonbreeding singing females, Figure 2B, comparison 5) was an exception (Supplementary Table 1). In this comparison, the spontaneously singing female canaries expressed a high number of Z chromosomal genes (260 genes, approximately 24% of all Z chromosomal genes) at higher levels than the breeding singing male canaries, which eliminated the male enrichment of Z chromosomal genes.

Sex-biased genes were not only Z chromosomal genes; in fact, sizeable numbers of the autosomal genes were found to be sex-biased genes, particularly female-biased genes (male-biased genes: >40%; female-biased genes: >96%, Figure 2B). Many studies have reported sex differences in the gene expression levels in multiple tissues and animal species (Frésard et al., 2013; Itoh et al., 2007; Nätt et al., 2014; Uebbing et al., 2015; Wolf and Bryk, 2011; Yang et al., 2006). Gene expression levels are generally under tight regulation, and perturbed expression levels might result in functional consequences. For example, several genes, including transcription factors, modulate distinct gene sets depending on their expression levels (Birchler et al., 2001; Doghman et al., 2013; Schulz, 2017).

We compared the male-biased genes in a pairwise manner across the comparisons and found that the male-biased autosomal genes showed low similarity to each other (Supplementary Figure 5A), whereas the male-biased Z chromosomal genes showed some similarity across the groups (Supplementary Figure 5B). Pairwise comparison of the female-biased genes across groups showed low similarity, and this finding was obtained for both autosomal genes (Supplementary Figure 5C) and Z chromosomal genes (Supplementary Figure 5D). Fisher’s exact test showed that chromosome 2 (comparisons 1, 4, and, 5) and chromosome 3 (comparisons 1, 2, and 4) are hotspots for female-biased genes; the enrichments of these chromosomes were observed in four out of the seven pairwise comparisons (Supplementary Table 1).

Further examination of the sex-biased genes (comparisons 1, 2, and 5) identified in the groups of naturally singing birds revealed that almost none of the male-biased autosomal genes (Figure 3A) and none of the female-biased Z chromosomal genes (Figure 3B) were shared across the three songbird species. In addition, relatively low numbers of male-biased Z chromosomal genes (Figure 3A) and female-biased autosomal genes (Figure 3B) were shared across the three songbird species. This observation indicates that only a small set of genes were regulated in the same manner (male- or female-biased) between species even though all birds exhibit singing behaviours. Moreover, a Gene Ontology (GO) enrichment analysis suggested that male- and female-biased genes did not functionally converge to similar pathways in the three studied species (Figure 3C).

**Figure 3.**
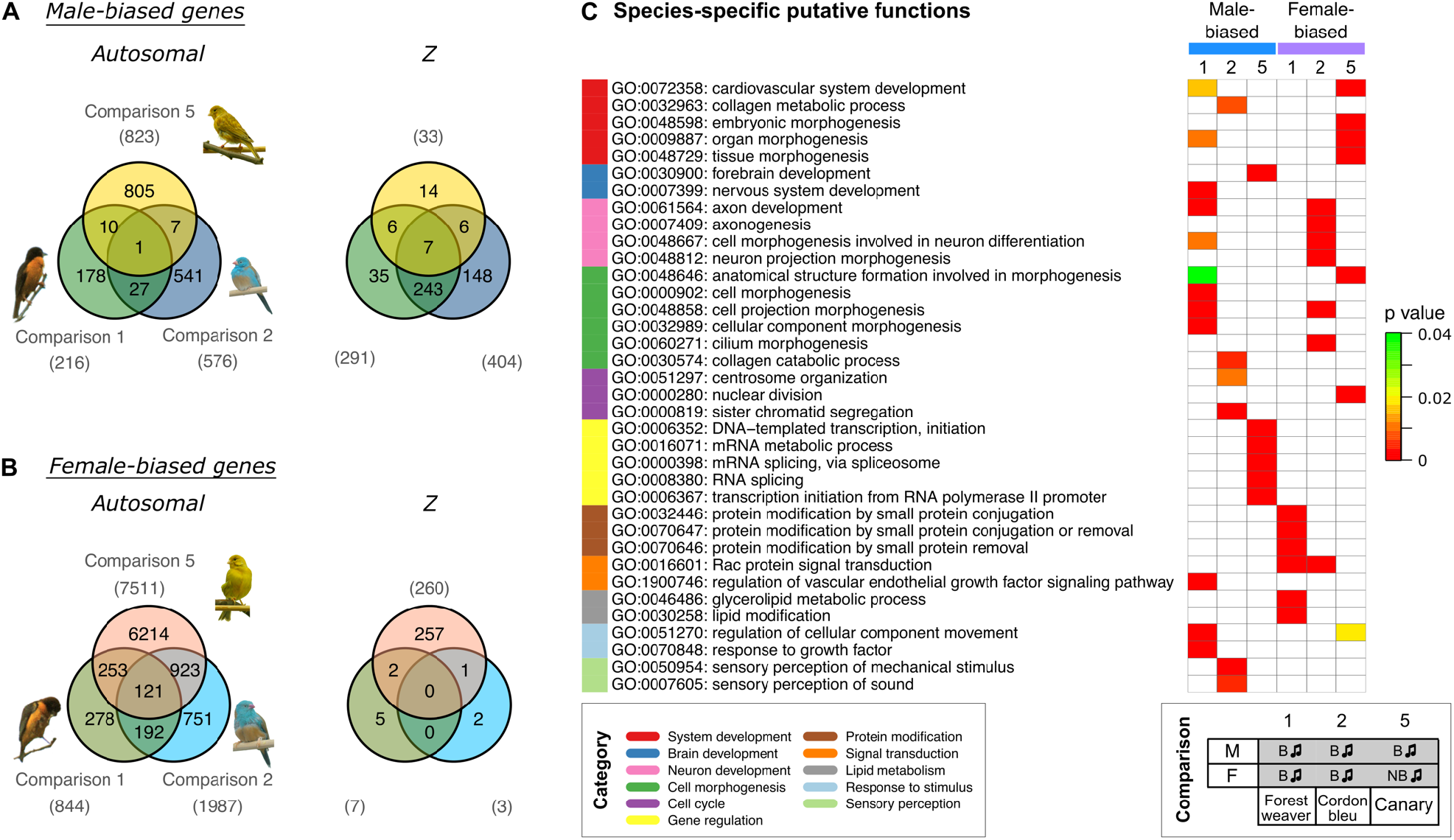
Cross-species comparisons of sex-biased genes expressed in the HVC of birds with singing phenotypes shows high species specificity. Venn diagrams of male-biased (**A**) and female-biased (**B**) autosomal and Z chromosomal genes. The numbers of sex-biased genes are indicated in parentheses. **C**, GO term enrichment analysis predicting the biological functions of sex-biased (autosomal and Z chromosomal) genes. The GO terms were categorized and are colour coded. Bonferroni-adjusted P values are shown by colour scales in the heatmap. Only five GO terms with the lowest Bonferroni-adjusted P values are shown for each set (see Supplementary Table 2 for the complete results). The phenotypes of the groups used for comparisons 1, 2, and 5 are listed at the bottom. Abbreviations: B, breeding; NB, nonbreeding. The music note indicates the presence or absence of singing behaviour

### The plasma androgen levels substantially alter the HVC transcriptomes of canaries

With the aim of understanding the features that distinguish the HVC transcriptomes in within-species contexts, we performed a principal component analysis (PCA) of the HVC transcriptomic data from the seven canary groups to identify variables that would explain the most variation in the data. By calculating the correlation coefficient between the PCs and the variables (plasma testosterone levels (Supplementary Figure 1), HVC volume (Figure 1A), sex, and singing), we identified the variables that were highly correlated with the most important PCs. PC1 explained 32% of the data variance (Figure 1C and Supplementary Figure 6) and was strongly correlated with the blood plasma androgen concentrations and the presence of singing activity (plasma androgen level: Pearson’s r = -0.57, Bonferroni-adjusted p = 1.0 × 10-4; singing: Pearson’s r = -0.51, Bonferroni-adjusted p = 0.001, Supplementary Table 3). Sex and HVC volume were strongly correlated with PC2 (sex: Pearson’s r = -0.91, Bonferroni-adjusted p = 4.4 × 10-16; HVC volume: Pearson’s r = -0.73, Bonferroni-adjusted p = 2.1 × 10-7), which explained 19% of the variance in the data (Supplementary Table 3). Taken together, the hierarchical clustering and PCA results suggest that although phylogenetic relationships dominate the variation between songbird species, the circulating testosterone levels, the presence of singing activity, and sex identity dominate the HVC gene expression patterns within a single species.

### The application of testosterone does not yield an HVC transcriptome that mimics that of natural singing canaries

To investigate the molecular mechanisms underlying the singing phenotype, we performed a differential gene expression analysis of the HVC transcriptomes of the two groups of singing canary females (nonbreeding spontaneously singing females and nonbreeding testosterone-stimulated singing females) against the nonsinging nonbreeding female canaries. Similarly, a differential gene expression analysis of the HVC transcriptomes of the two groups of singing male canaries (breeding singing males and nonbreeding testosterone-stimulated singing males) with nonsinging nonbreeding male canaries was performed. A substantial number of genes were differentially expressed between singing and nonsinging birds (Figure 4A, nonbreeding singing females: 4,125 genes; nonbreeding testosterone-stimulated singing females: 7,702 genes; breeding singing males: 3,359 genes; nonbreeding testosterone-stimulated singing males: 5,106 genes). Approximately 65% of the differentially expressed genes found in the naturally singing birds overlapped with those found in the testosterone-treated birds of the same sex (male: 65%; female: 62%). However, both testosterone-treated groups had markedly higher numbers of differentially expressed genes than the naturally singing groups of the same sex (Figure 4A). Thus, although testosterone implantation induced singing in females and males, most testosterone-responsive genes (female: 67%; male: 57%) might not be necessary for singing behaviour *per se* but rather a response to nonphysiological levels of testosterone (Supplementary Figure 1). Alternatively, mechanisms that lead to the first song in life might be very different from those that reinduce singing in animals that sang before (Vellema et al., 2019a). Because male canaries sing regularly starting from approximately 50 days of age (Nottebohm et al., 1986) while most female canaries never sing (Ko et al., 2020), the induction of singing in females by testosterone likely activated genes related to first-time singing, whereas in males, this treatment activated genes related to reinduced singing.

**Figure 4.**
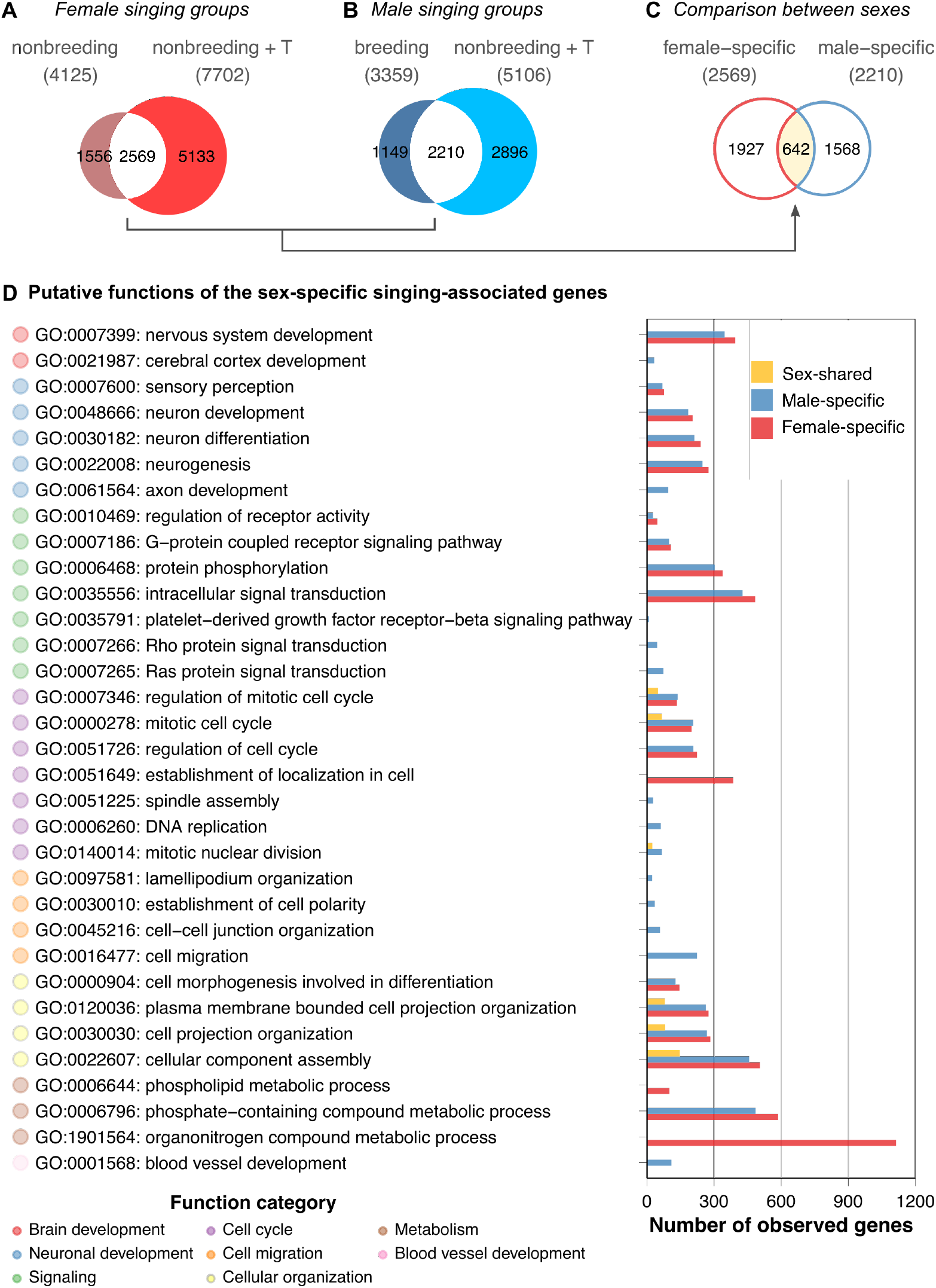
HVC transcriptomes of naturally singing canaries are dissimilar from testosterone-stimulated singing canaries. Venn diagrams comparing (**A**) the two female singing groups, (**B**) the two male singing groups, and (**C**) the female- and male-specific expressed genes resulting from comparisons A and B. Note that only approximately 20% of the genes were shared among the singing females and approximately 26% were shared among the singing males. The majority of these overlapping genes were not sex-shared; approximately 75% of female intersections were specific to female groups, whereas 71% of male intersections were specific to male groups. **D**, GO term enrichment analysis of female-specific, male-specific, and sex-shared expressed genes derived from comparisons A to C. Because many putative functions show similarities between male- and female-specific expressed genes, the results suggest functional intraspecies convergence based on sex-specific gene expression in the HVC. See Supplementary Table 7 for the full results of the GO term enrichment analysis.

### The majority of differentially expressed genes between singing and nonsinging canaries are sex-specific

We examined whether the overlapping gene sets of the female canary signing groups were similar to the overlapping gene sets of the singing males (Figure 4C). The Venn diagram indicated that only approximately 25% (642 of 2569 genes, Supplementary Table 4) of the female-specific expressed genes and 29% (642 of 2210 genes, Supplementary Table 5) of the male-specific expressed genes were shared by both sexes of singing canaries (Supplementary Table 6).

### The sex-specific genes show functional overlap

To understand the putative biological functions of the female-specific, male-specific, and sex-shared expressed genes in the HVC, we performed a GO term enrichment analysis. Interestingly, the results suggested that the female- and male-specific expressed genes largely overlapped at the functional level (Figure 4D and Supplementary Table 7). GO terms such as nervous system development (GO:0007399), neuron development (GO:0048666), and intracellular signal transduction (GO:0035556) were shared among the sexes. Female-specific GO terms were mainly related to cellular maintenance, such as organonitrogen compound metabolic process (GO:1901564) and phospholipid metabolic process (GO:0006644). In contrast, the male-specific GO terms were axon development (GO:0061564), DNA replication (GO:0006260), cell migration (GO:0016477), and blood vessel development (GO:0001568). In summary, although only approximately one-fourth to one-third of the sex-specific expressed genes are shared among the sexes in terms of their identities, most of the predicted functions of the sex-specific expressed genes are nevertheless sex-shared in singing canaries.

All four groups of singing canaries had elevated plasma androgen levels compared with nonbreeding nonsinging canaries of the same sex (Supplementary Figure 1 and (Ko et al., 2020)). The activation of singing is likely testosterone-dependent in females as in males (Hartley and Suthers, 1989; Heid et al., 1985; Leitner et al., 2001a; Nottebohm et al., 1987). Thus, the potential master regulator for inducing singing behaviour might be testosterone-sensitive and Z-linked. One such candidate is *DMRT1* (doublesex and mab-3 related transcription factor 1), which is present in the sex-shared expressed gene list. The Z-linked gene *DMRT1* is needed for male sex determination in birds and other animal species (Herpin and Schartl, 2015; Lambeth et al., 2014; Smith et al., 2009). The overexpression of *DMRT1* in female chicken embryos reduces aromatase expression in the gonads and triggers development of the testis (Lambeth et al., 2014). The role of DMRT1 in adult avian tissues in general and in the brain in particular is unknown. Thus, whether DMRT1 affects steroid metabolism in the HVC, such as converting testosterone to more active metabolites (see below), or whether it regulates other mechanisms that direct the HVC into a configuration that enables singing needs to be validated by future experiments.

Testosterone can be converted to 5α-dihydrotestosterone (5α-DHT) and 17β-estradiol, which activate androgen receptor (AR) and estrogen receptors (ERα and ERβ), respectively. Both AR (encoded by *AR*) and ERs (ERα encoded by *ESR1* and ERβ encoded by *ESR2*) are transcription factors that play important roles in the transcription of numerous genes (Bourdeau et al., 2004; Pihlajamaa et al., 2015; Takayama et al., 2007; Wilson et al., 2016). Interestingly, *ESR1* was specifically expressed in males, whereas *ESR2* was female-specifically expressed in the HVC of canaries, which suggests that estrogen receptor paralogues could provide finer-tuned mechanisms for sex-specific regulation. Empirical results have shown that ERα and ERβ bind to the same estrogen response element motifs (Zhao et al., 2010) and might functionally overlap in some tissues. Moreover, ER paralogues show sex differences in expression levels and tissue specificity (Zhang et al., 2017). In quail, the administration of an agonist specific to ERβ on embryonic day 7 demasculinizes male sexual behaviour and midbrain nuclei characteristics in Japanese quails, whereas an agonist specific to ERα does not exert this effect (Court et al., 2020). The specific roles of each ER paralogue in the adult HVC of male and female songbirds warrant further investigation. Until now, ERα but not ERβ was expected to regulate the function of HVC neurons of adult songbirds in addition to the AR (Frankl-Vilches and Gahr, 2017).

## Conclusion

In this study, we investigated the sex differences in the gene expression patterns in the HVC of three songbird species with different levels of sex-specific singing and several different song-related phenotypes between male and female canaries. Our inter- and intraspecies comparisons yielded large-scale transcriptional sex differences regardless of singing behaviour. Instead, fundamental sex differences in gene expression levels were found in the HVC, and these differences were highly species-specific. By leveraging several experimental groups of canaries, we found that the plasma androgen levels and sex were the major contributors to the variations in the HVC transcriptome. Although testosterone reliably induced singing in both female and male canaries, testosterone treatment did not alter the transcriptome to imitate that of natural singing birds. Our results suggest that female and male canaries rely on different gene networks for singing behaviour, but the sex-specific gene networks might show functional convergence.

## Data accessibility

The microarray CEL files (GEO Series accession number <u>GSE83674)</u> are available on NCBI’s Gene Expression Omnibus. The processed microarray data, plasma androgen levels, HVC volume measures, and all scripts used for analysis and visualization are available in GitHub (https://github.com/maggieMCKO/SongbirdSexDiff).

## Authors’ contributions

M-CK collected the nonbreeding testosterone-implanted canaries, analysed and visualized the transcriptomic data and was a major contributor to the writing of the manuscript. AB and M-CK performed the RNA extractions and microarray hybridizations. MG and M-CK isolated the tissues for microarray. MG collected the forest weavers and cordon-bleus. MG conceived the study and obtained financial support for the study. M-CK, CF-V and MG drafted the manuscript. All the authors read and approved the final manuscript.

## Acknowledgements

We thank Dieter Schmidl for catching the forest weavers, Stefan Leitner for providing the breeding canaries, Roswitha Brighton and David Witkowski for maintaining the breeding colonies of the blue-capped cordon-bleus and the canaries, Doris Walcher for providing data from the breeding male and female canaries as well as the nonbreeding nonsinging male and female canaries, and Wolfgang Goymann and Monika Trappschuh for the RIA of the plasma testosterone. Furthermore, we express our appreciation to Maude Baldwin, Glenn Cockburn, Falk Dittrich, Lisa Gill, Vincent Van Meir, Keren Sadanandan, and Michiel Vellema for their comments on previous versions of this manuscript and to the International Max Planck Research School for Organismal Biology for the training and support provided.

## Tables

**Supplementary Table 1. Fisher’s exact tests of sex-biased genes for chromosome enrichment**.

**Supplementary Table 2. GO term enrichment analysis of sex-biased genes identified from the forest weaver (comparison 1), cordon-bleu (comparison 2), and canary (comparison 5) comparisons**.

Abbreviations: FWm, forest weaver male-biased genes; CBm, cordon-bleu male-biased genes; Cm, canary male-biased genes; FWf, forest weaver female-biased genes; CBm, cordon-bleu female-biased genes; CfS, canary female-biased genes.

**Supplementary Table 3. Pearson’s correlation analysis of principal components and variables (plasma androgen levels, HVC volume, sex, and singing)**.

**Supplementary Table 4. Female-specific expressed genes**.

Abbreviations: SDfS: nonbreeding spontaneously singing female canaries; SDfT: nonbreeding testosterone-stimulated singing female canaries.

**Supplementary Table 5. Male-specific expressed genes**.

Abbreviations: LDm: breeding singing male canaries; SDmT: nonbreeding testosterone-stimulated singing male canaries.

**Supplementary Table 6. Sex-shared expressed genes**.

Abbreviations: LDm: breeding singing male canaries; SDfS: nonbreeding spontaneously singing female canaries; SDmT: nonbreeding testosterone-stimulated singing male canaries; SDfT: nonbreeding testosterone-stimulated singing female canaries.

**Supplementary Table 7. GO term enrichment analysis of female-specific, male-specific and sex-shared genes**.

## Supplementary Figures

**Supplementary Figure 1.**
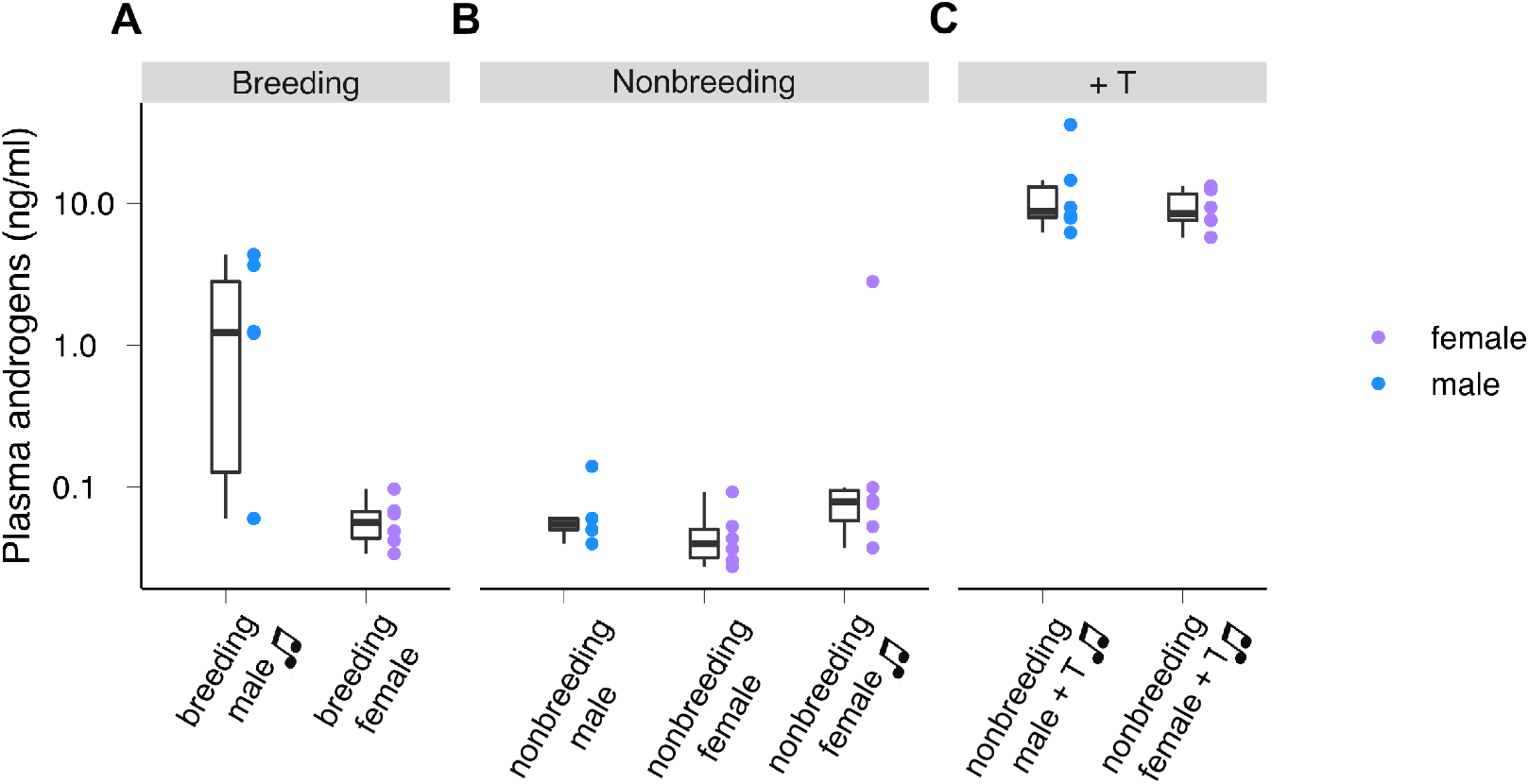
Plasma androgen levels of seven groups of canaries on the day of sacrifice. **A**, Canaries in breeding conditions. Singing males 1.76 ng/ml (mean), nonsinging females 59.2 pg/ml. Mann-Whitney Test, U = 6, P value = 0.07. **B**, Canaries in nonbreeding conditions. Nonsinging males 66.7 pg/ml (mean), nonsinging females 47.2 pg/ml, singing females 52.6 pg/ml. Mann-Whitney Test (nonsinging males vs. nonsinging females), U = 9, P value = 0.172. Mann-Whitney Test (nonsinging males vs. singing females), U = 24, P value = 0.377. **C**, Testosterone-implanted nonbreeding canaries. Males 13.5 ng/ml, females 9.31 ng/ml. Mann-Whitney Test, U = 13, P value = 0.485. Testosterone implantation significantly increased the plasma androgen levels of both nonbreeding males and females Mann-Whitney Test (nonbreeding nonsinging males vs. testosterone-implanted singing males), U = 0, P value = 0.00492. Mann-Whitney Test (nonbreeding nonsinging females vs. testosterone-implanted singing females), U = 0, P value = 0.00217. The plasma androgen levels were higher in the breeding males than in the nonbreeding nonsinging males (Mann-Whitney Test, U = 32, P value = 0.0275). The boxes indicate the 25th/50th/75th percentiles (bottom/middle/top bar), and the extent of the whiskers indicates the most extreme values that are within 1.5 times the IQR (interquartile range) of the hinge. Each colour-coded dot indicates the measurement from one bird. The nonbreeding singing female canaries data were obtained from (Ko et al., 2020).

**Supplementary Figure 2.**
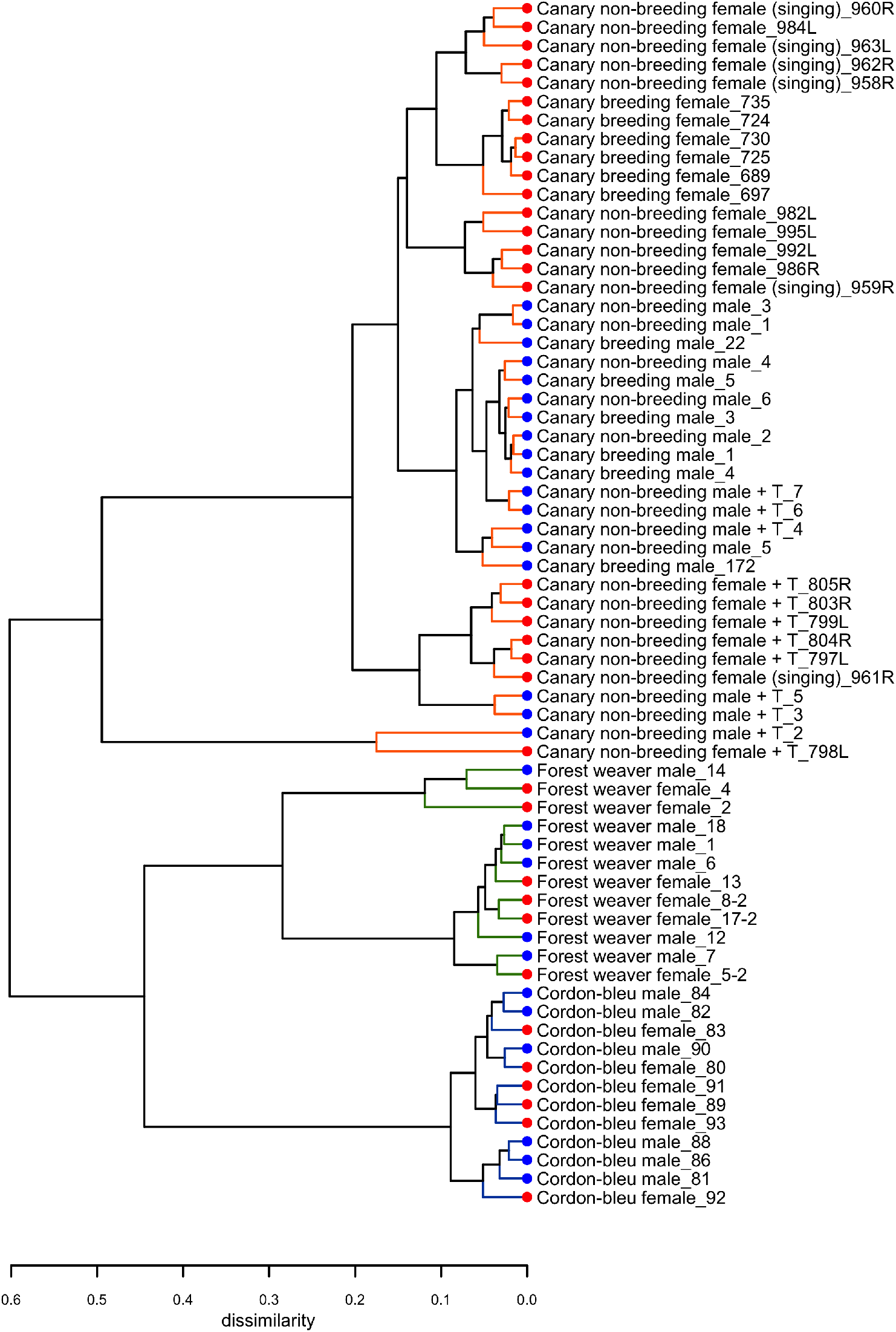
Hierarchical clustering of the HVC transcriptomes of 65 birds used in this study. Hierarchical clustering showed that the HVC transcriptomes were first clustered based on the phylogenetic relationship; among canaries, nonbreeding females implanted with testosterone showed the most distinctive patterns. +T: testosterone implantation.

**Supplementary Figure 3.**
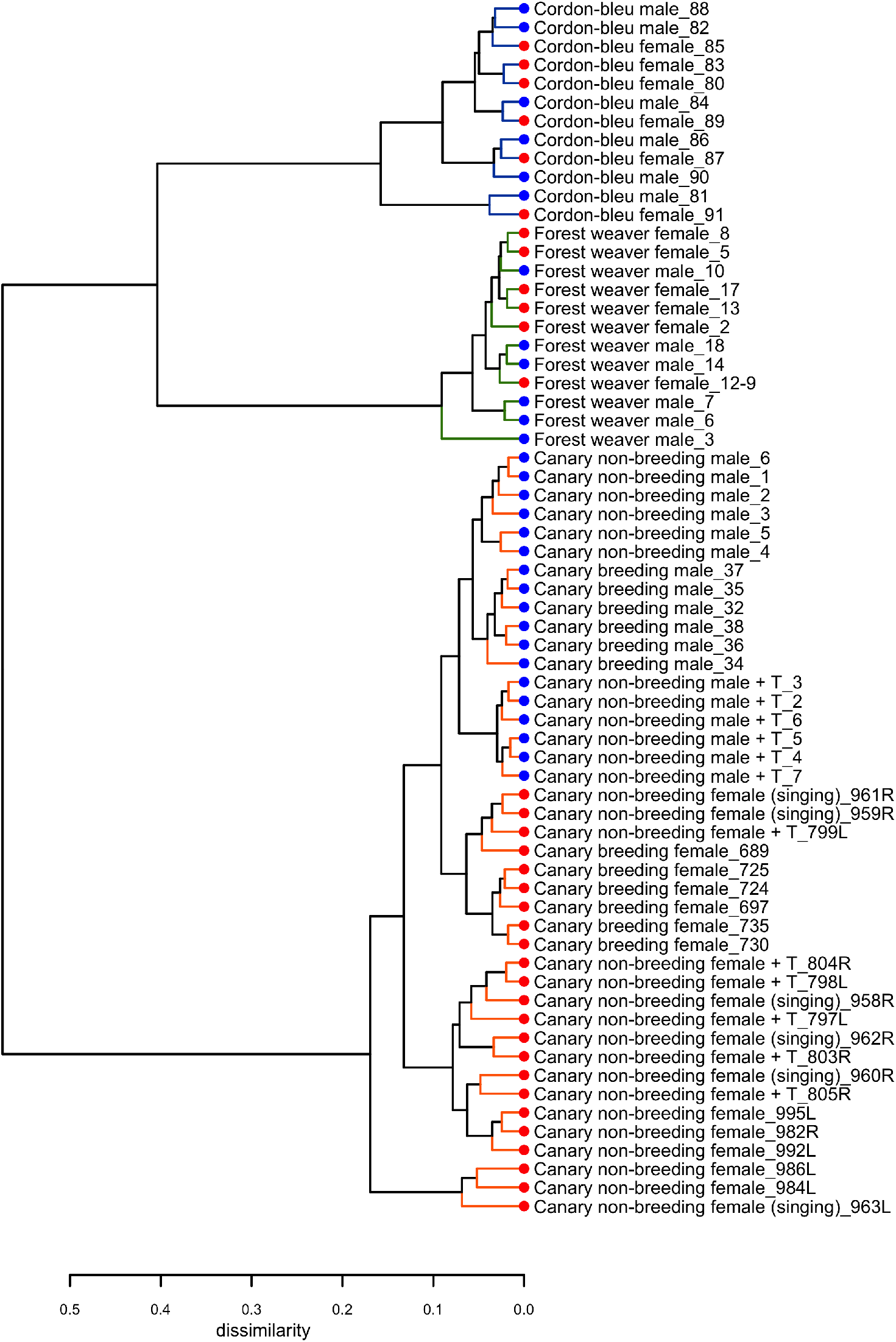
Hierarchical clustering of the entopallium transcriptomes of 65 birds used in this study. Hierarchical clustering showed that the entopallium transcriptomes were first clustered based on the phylogenetic relationship. The canaries were clustered by sex. +T: testosterone implantation.

**Supplementary Figure 4.**
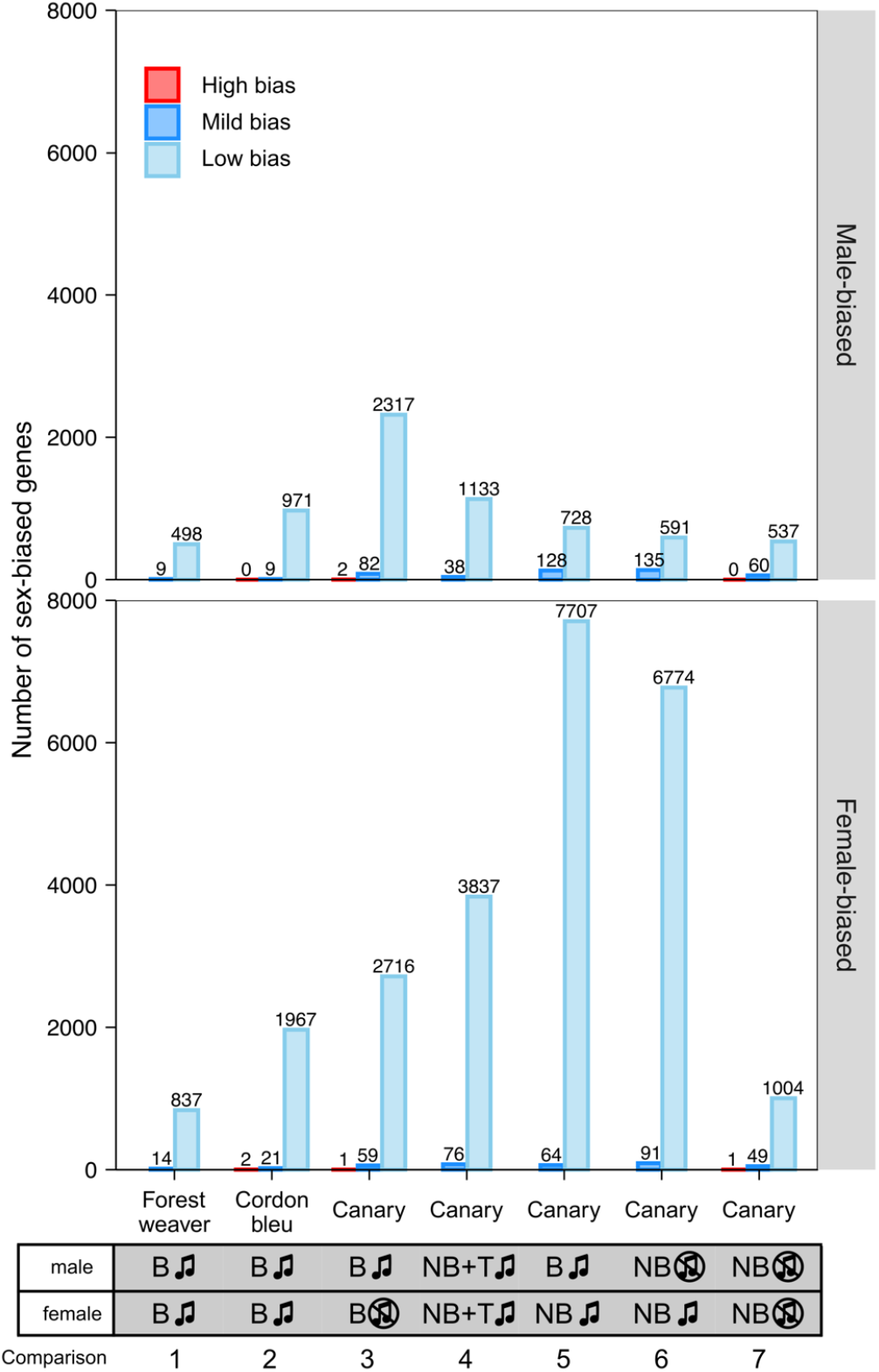
The majority of sex-biased genes showed low sex bias. The bar graph summarizes the number of sex-biased genes in the HVC transcriptome identified from each male-to-female comparison. High bias: |log_2_(fold change)| ≥ 2; moderate bias: 1 ≤ |log_2_(fold change)| < 2; low bias: 0.5 ≤ |log_2_(fold change)| < 1. The phenotypes of the groups being compared in each male-to-female comparison are listed at the bottom of the graph.

**Supplementary Figure 5.**
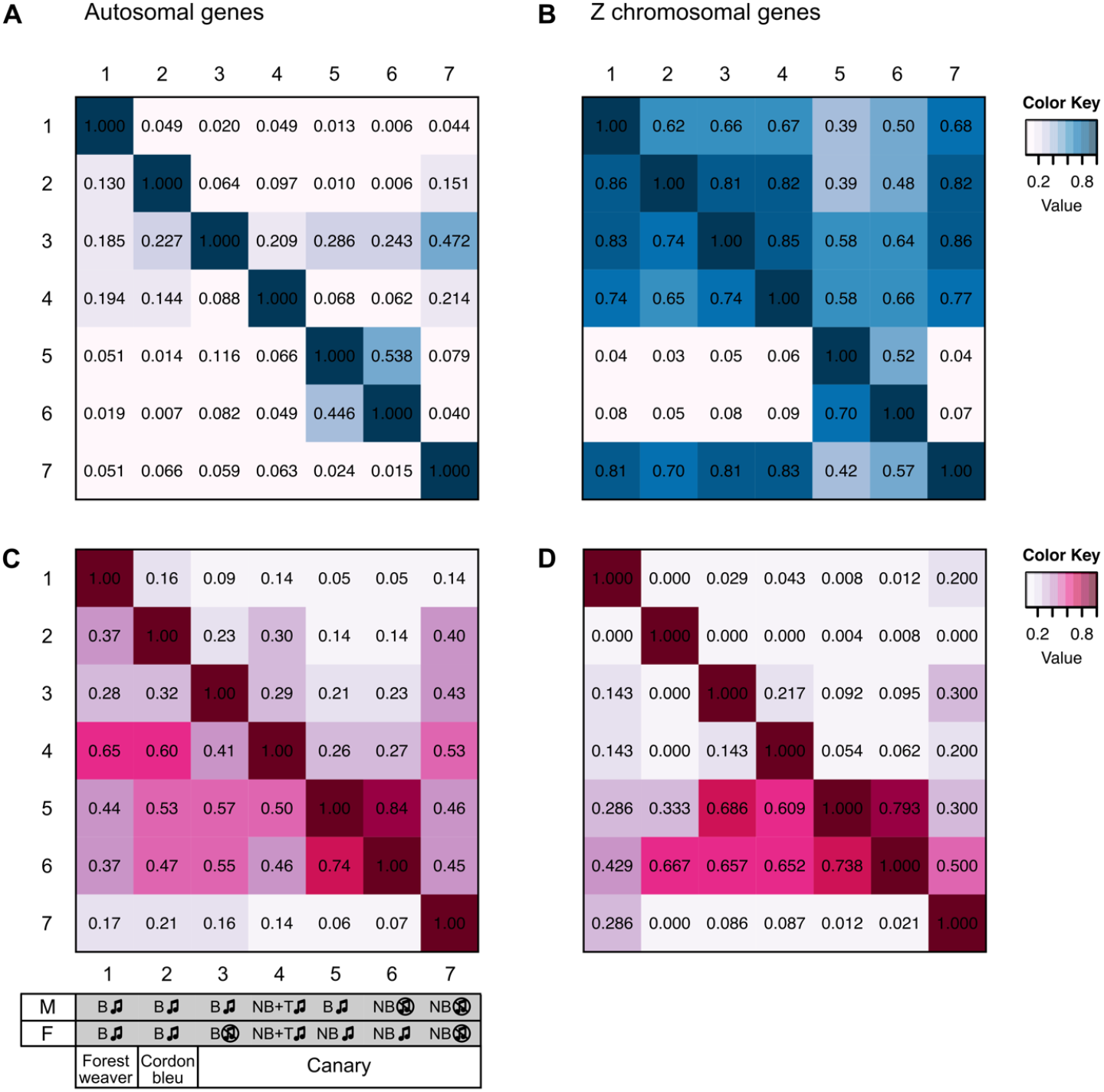
Pairwise comparisons between autosomal and Z chromosomal sex-biased genes. **A**, Autosomal male-biased genes. **B**, Z chromosomal male-biased genes. **C**, Autosomal female-biased genes. **D**, Z chromosomal female-biased genes. The numbers indicated in the matrices are the proportions of identical genes identified from each pair. The phenotypes of the groups used for each comparison are listed at the bottom. B: breeding; NB: nonbreeding; T: testosterone implantation; the music note indicates the presence or absence of singing behaviour.

**Supplementary Figure 6.**
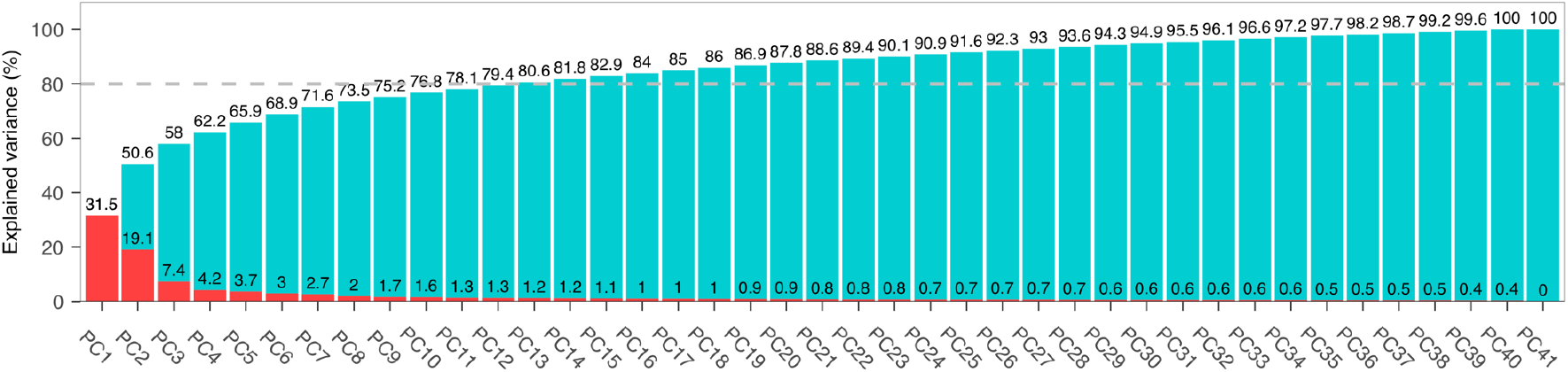
PCA scree plot showing the percentage of variance explained by each principal component.

